# A simple pressure-assisted method for MicroED specimen preparation

**DOI:** 10.1101/665448

**Authors:** Jingjing Zhao, Hongyi Xu, Hugo Lebrette, Marta Carroni, Helena Taberman, Martin Högbom, Xiaodong Zou

## Abstract

Micro-crystal electron diffraction (MicroED) has shown great potential for structure determination of macromolecular crystals too small for X-ray diffraction. However, specimen preparation remains a major bottleneck. Here, we report a simple method for preparing MicroED specimens, named Preassis, in which excess liquid is removed through an EM grid with the assistance of pressure. We show the ice thicknesses can be controlled by tuning the pressure in combination with EM grids with appropriate hole sizes. Importantly, Preassis can handle a wide range of protein crystals grown in various buffer conditions including those with high viscosity, as well as samples with low crystal contents. Preassis is a simple and universal method for MicroED specimen preparation, and will significantly broaden the applications of MicroED.

## Introduction

Three-dimensional electron diffraction, known as micro-crystal electron diffraction (MicroED)(Shi *et al*., 2013; Nannenga *et al*., 2014), has shown great potential for structure determination of macromolecules from nano- and micron-sized crystals which are too small for X-ray diffraction(Xu *et al*., 2019; Nannenga & Gonen, 2019; Clabbers & Xu, 2020). This method has attracted large interests and extensive attention in structural biology, structural chemistry, and the pharmaceutical industry during the past years. However, until now, only 16 protein structures solved by MicroED are reported in the PDB database, all except R2lox(Xu *et al*., 2019) had been previously determined by X-ray diffraction. The R2lox structure(Xu *et al*., 2019) was solved by molecular replacement using a search model of 35% sequence identity (PDB code 6QRZ). MicroED data collection has been very fast (1-3 minutes)(Nannenga *et al*., 2014; Gemmi *et al*., 2015; Nederlof *et al*., 2013) by applying continuous rotation of crystals and a hybrid pixel detector or complementary metal-oxide semiconductor (CMOS) detector in movie mode. Data processing and structure determination can be done using standard X-ray crystallographic software suites. Despite those advances in MicroED experiments, the major bottleneck has been specimen preparation, which is most delicate and time-consuming.

In MicroED experiments, sub-micrometer thick crystals are needed in order to allow the electron beam to penetrate through the specimen and minimize multiple scattering. The surrounding ice also needs to be as thin as possible to improve the signal-to-noise ratio while still protecting the protein crystals from dehydration. Suitable crystal size can be obtained by adjusting the crystallization conditions(Russo Krauss *et al*., 2013; McPherson & Gavira, 2014), by segmenting large crystals using mechanical forces (e.g. vigorous pipetting, sonication, vortexing with beads)(de la Cruz *et al*., 2017), or by focused ion beam milling under cryogenic conditions (cryo-FIB)(Duyvesteyn *et al*., 2018; Zhou *et al*., 2019; Martynowycz *et al*., 2019; Beale *et al*., 2020). High molecular weight polymers, like polyethylene glycols (PEG), are common and popular agents to produce volume-exclusion effects for successful protein crystallization(McPherson & Gavira, 2014; Russo Krauss *et al*., 2013; Abdallah *et al*., 2016). However, their addition makes the buffer viscous. Lipid cubic phase (LCP) is commonly used for crystallization of membrane protein crystals, and they are extremely viscous as toothpaste(Martynowycz *et al*., 2019; Caffrey, 2015; Polovinkin *et al*., 2020). It has been very challenging to prepare MicroED samples for crystals grown in viscous buffers(Nannenga & Gonen, 2019; Martynowycz *et al*., 2019; Shi *et al*., 2016*a*). The pipetting-blotting-plunging rountine(Dubochet & McDowall, 1981), originally designed for single-particle cryoEM specimen preparation, has two major drawbacks for MicroED specimen preparation; 1) many microcrystals are removed by blotting and 2) it is insufficient in removing viscous liquids(Shi *et al*., 2016*b*). Manual back-side blotting(Shi *et al*., 2016*b*), direct crystallization on EM grids(Li *et al*., 2018), and cryo-FIB(Martynowycz *et al*., 2019; Beale *et al*., 2020; Polovinkin *et al*., 2020; Martynowycz *et al*., 2020) have been proposed as possible solutions to deal with the above problems. However, a simple and universal method for MicroED specimen preparation is still missing. There is an urgent need to develop such a method in order to apply MicroED on a wide range of macromolecular crystals.

Here, we describe a pressure-assisted method, named Preassis, for the preparation of MicroED specimens. The excess liquid is removed through the EM grid with the assistance of pressure. Preassis is applicable for a wide range of protein crystal suspensions with both low and high viscosities, and can preserve 10 times more crystals on the TEM grid compared with Vitrobot. The ice thicknesses can be controlled by tuning the pressure in combination with EM grids with appropriate hole sizes. We provide detailed experimental guidance in finding appropriate parameters for preparing MicroED specimens of new protein crystal samples. More importantly, the Preassis method is simple and easy to implement, making it widely accessible to cryoEM labs at a very low cost.

## Results and discussions

The basic concept of Preassis is to pull a portion of sample suspension through an EM grid and simultaneously remove excess liquid from the backside of the grid with the assistance of suction/pressure. In its simplest setup (**Fig. 1a**), an EM grid is placed on a filter paper which is rested on the mouth of a Buchner flask and pumped with a certain pumping speed. A droplet of the sample suspension is then deposited onto the grid. Due to the suction underneath the filter paper, the excess liquid is “pulled” through the grid. Then the grid is manually picked up using a tweezer and plunged into liquid ethane. Details of this setup and specimen preparation procedures are available in the **Supplementary Protocol**. The overall vitrified ice thickness on the EM grid can be tuned by changing the pressure, hole size of the EM grid, and the time over which the pressure is applied. While the pressure can be changed continuously, the change of the hole size is done by choosing the type of EM grids. In the setup described above, the pressure is adjusted by changing the pumping speed, which is linearly proportional to the pumping speed in the range of 20% to 80% (**Supplementary Fig. 1**). It is worth noting that Preassis can be applied on pre-clipped EM grids used for auto-loading as well, which makes this method very promising for future automation.

**Fig 1.**
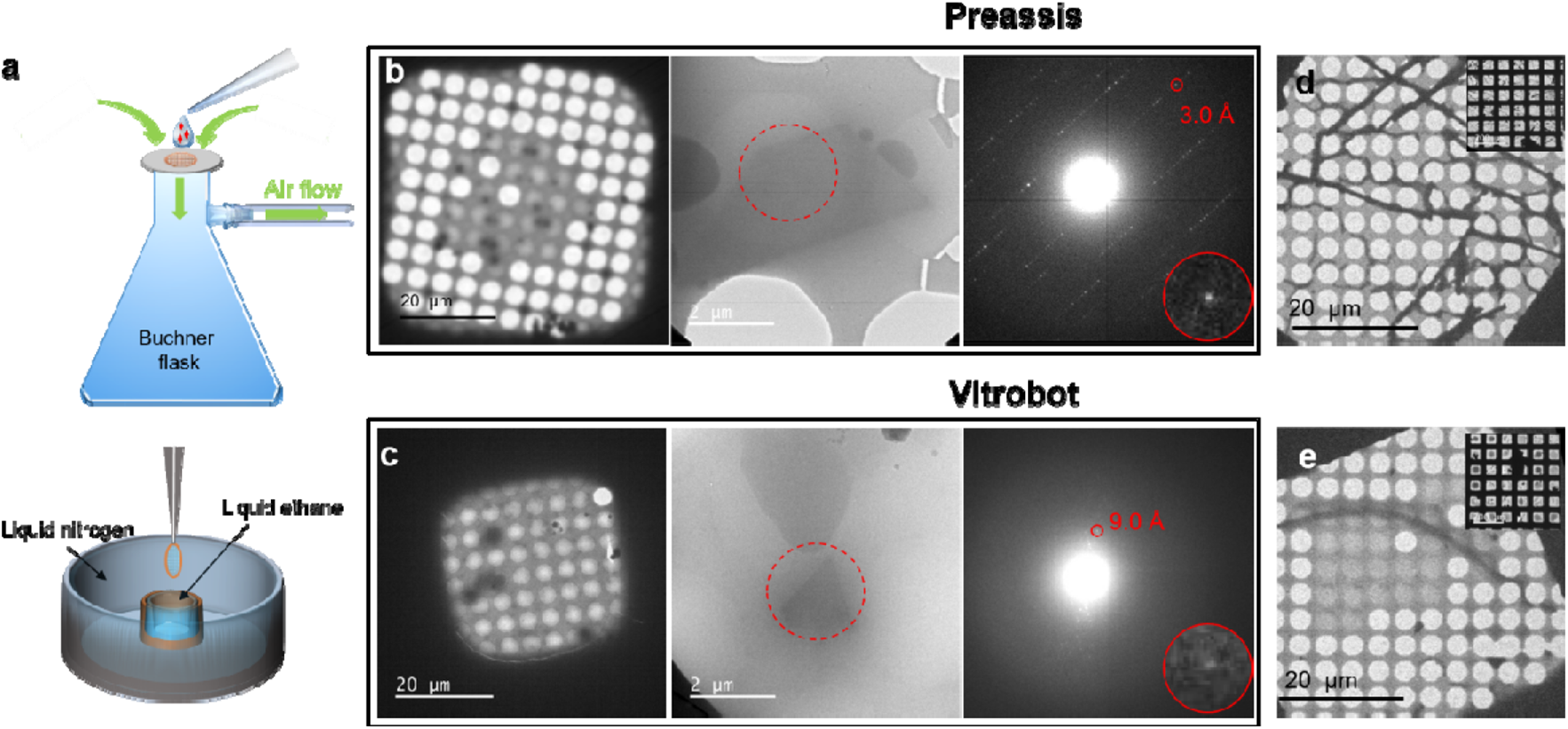
Schematic drawing of the Preassis setup and a comparison of specimen preparation between Preassis and Vitrobot. **a**, A drop of the microcrystal suspension is transferred onto an EM grid rested on a support (e.g. filter paper). Simultaneously, the extra liquid is removed through the support with the assistance of pressure, and followed by a manually plunge freezing in liquid ethane for vitrification. **b**-**c**, TEM images and diffraction patterns taken from specimens of viscous R2lox suspensions prepared by Preassis (**b**) and Vitrobot (**c**). The R2lox crystals were grown in a highly viscous mother liquid with 44% PEG 400. The resolution of electron diffraction pattern was improved from 9.0 Å to 3.0 Å by Preassis. **d-e**, TEM images taken from specimens prepared by Preassis (**d**) and Vitrobot (**e**), using a orthorhombic lysozyme crystal sample. It shows that Preassis can preserve 10 times more crystals on the grid than Vitrobot can.

The most important advantage of Preassis is its ability to handle protein crystals grown in highly viscous buffers. We applied it to crystals of *Sulfolobus acidocaldarius* R2-like ligand-binding oxidase (R2lox) (**Fig. 1b** and **c**) grown with 44% PEG 400. It was difficult to remove enough amount of liquid by Vitrobot even with extreme blotting conditions (2 layers of filter paper on each side, strong blotting force (16), and long blotting time (10 s)), as shown in Fig. 1c. Only a few grid squares were electron beam transparent and the ice layers are very thick (as represented in **Supplementary Fig. 2b** and **c**), making it extremely difficult to obtain sufficient MicroED data with good diffraction quality. The diffraction quality of MicroED data of R2lox couldn’t be improved and the R2lox project got stuck for more than one year until the Preassis method was proposed. Using Preassis with 30.7 mbar pressure and Quantifoil grid R3.5/1, the viscous liquid was efficiently removed obtaining a lot of grid squares suitable for searching crystals for MicroED data collection, as shown in **Fig. 1b**. Owing to the reduced vitrified ice thickness, the resolution of electron diffraction (ED) data was significantly improved from 9.0 Å to 3.0 Å (as shown by ED patterns in **Fig. 1b** and **c**). Consequently, sufficient and high quality MicroED data were collected from one single TEM grid prepared by Preassis, and a typical MicroED dataset of R2lox is shown in **Supplementary Video 1**. Preassis is crucial for the successful structure determination of R2lox, the first novel protein structure solved by MicroED(Xu *et al*., 2019). Furthermore, Preassis can preserve 10 times more crystals on the EM grid than that of Vitrobot, as shown in **Fig. 1d** and **e**. This is mainly because that Preassis takes away the excess liquid from the back-side (copper-side) of an EM grid and the holey carbon layer of the grid can act as a sieve to keep more crystals, while the Vitrobot uses two-side blotting and the excess liquid and precious crystals can be taken away from both sides.

It is worth noting that unlike single particle cryo-EM experiments where 10 000 to 100 000 of particles are required for a complete structure determination, MicroED experiments require only a limited number (up to 50) of crystals. Therefore, a successful MicroED specimen preparation is to ensure there are sufficient microcrystals covered by thin ice on several grid squares, so that high-quality MicroED data can be collected. The influences of pressure and hole sizes were investigated by combining TEM images and SAED patterns. Each specimen preparation condition was repeated 2 to 5 times and the reproducibility is relatively good, as illustrated in **Supplementary Fig. 3**. It is worth mentioning that the ice layer thickness can vary throughout an EM grid prepared by the current Preassis setting, as shown in **Supplementary Fig. 4**. This may be due to non-uniform contact of the EM grid to the filter paper. Such a gradient of ice thickness may not be a disadvantage, and instead increase the chance of finding suitable crystals for MicroED data collection.

**Fig. 2** illustrates how the pressure and hole size affect the overall ice thickness on the EM grid and the resulting quality of electron diffraction patterns using orthorhombic lysozyme crystals as an example(Xu *et al*., 2018*a*). By applying a low pressure of 17.2 mbar in a combination of EM grids with a small hole size of 1.2 µm (R 1.2/1.3 Quantifoil grids), usable specimens could be obtained (**Fig. 2b**). However, the vitrified ice was relatively thick as seen by the reduced transparent area in each grid square and strong ice rings in corresponding ED patterns (**Fig. 2b**). A better EM grid with thinner ice layers could be obtained by slightly increasing the pressure to 27.7 mbar, where the sharp edges of grid squares are visible in the TEM images (**Fig. 2a**). With thinner ice, it was easier to find suitable crystals for MicroED data collection, and ice rings were eliminated in the ED patterns. Consequently, the quality of MicroED data was improved in terms of resolution and signal to noise ratio.

**Fig 2.**
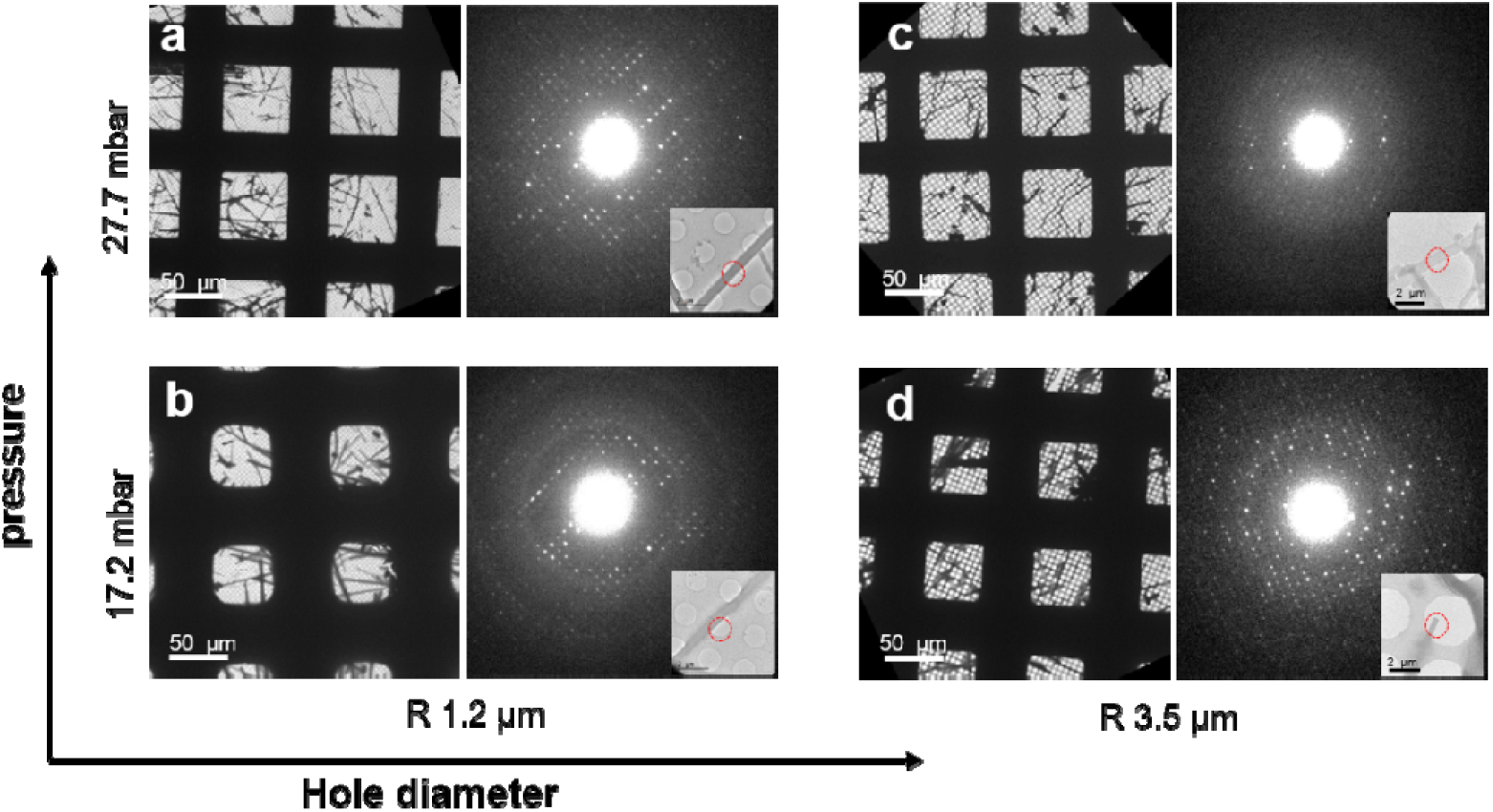
Adjustment of vitrified ice thickness by tunning pressure and choosing EM grids with different hole sizes. **a-d**, Representative low magnification TEM images and typical diffraction patterns taken from lysozyme crystal specimens prepared under different pressures (17.2 and 27.7 mbar) and Quantifoil grids with different hole sizes (R 1.2/1.3 and R 3.5/1). Insets display the areas of lysozyme crystals (marked by red circles) from which the diffraction patterns were taken. This comparison shows both the pressure and hole size of the grid play important roles in ice thickness adjustment.

Furthermore, the hole size of the EM grid has a strong influence on the ice thickness. Instead of increasing the pressure, thinner ice layers can also be obtained by using EM grids with larger holes because it is easier for liquids to pass through. When a R 3.5/1 grid with a hole diameter of 3.5 µm was used (**Fig. 2d**), the ice became thinner compared to that of R 1.2/1.3 grids prepared under the same pressure (17.2 mbar, **Fig. 2b**). High resolution reflections could be obtained in the corresponding ED pattern and no ice rings are observed. On the other hand, a combination of high pressure (27.7 mbar) with a large hole size (R 3.5/1 grid) led to slight dehydration of the lysozyme crystals, as indicated by reduced resolution in the ED pattern (**Fig. 2c**). Our results show that optimal ice layer thicknesses could be obtained under several conditions; both combinations with high pressure/small hole size (**Fig. 2a**) and low pressure/large hole size (**Fig. 2d**) produced optimal EM grids for MicroED data collection. This is significant because it allows us to select EM grids according to the size and shape of the crystals, and control and fine-tune the ice thickness by adjusting the pressure. Ideally, the hole size of the grid should be slightly smaller than or comparable to the largest crystal dimensions to maximize the hole areas and at the same time prevent the loss of crystals through the holes. This is particularly important when the initial density of crystals is low. We anticipate that the size and shape of microcrystals may also affect the ice thickness. Therefore Preassis was applied also to tetragonal lysozyme crystals with an isotropic shape and the results are shown in **Supplementary Figs. 4** and 5**a**. Compared to the optimal pressure (27.2 mbar) for the preparation of orthorhombic lysozyme crystals with rod-like morphology on the R 1.2/1.3 grids, a higher pressure (37.2 mbar) was needed to achieve similar ice thickness and MicroED data quality for the tetragonal lysozyme crystals. It is observed that despite the absence of vitrified ice in the neighbouring empty holes, most crystals are still embedded in ice, showing microcrystals attract more liquids than the EM grids do presumably caused by the stronger surface tension. Consequently, the applied pressure needs to be adjusted according to the size and shape of the crystals.

For non-viscous samples (e.g. no high molecular weight polymers in the buffer), usable specimens can be prepared by Preassis even without applying pressure if other parameters (e.g hole size and time) are carefully chosen (**Supplementary Fig. 5**). If the crystal suspension is viscous due to e.g. high concentration or high molecular weight of PEGs in the buffer, high pressures (e.g. > 180 mbar) are required. To systematically study the specimen preparation under different viscosities, several experiments were conducted with various concentrations of PEG 6000 mixed with microcrystals. Submicron-sized crystals of an inorganic sample zeolite ZSM-5 were used instead since it is difficult to obtain the same type of protein crystals from mother liquids of different viscosities. The results are shown in **Supplementary Fig. 6**. We found that, for crystal suspension with PEG 6000 lower than 25%, usable grids could be obtained without applying pressure when grids with large holes were used (e.g. Quantifoil R 3.5/1, **Supplementary Fig. 6f** and **g**). However, the amorphous ice on grids prepared under those conditions was relatively thick in most grid squares, which reduces the contrast of the crystals and makes it difficult to find suitable crystals for MicroED data collection. By applying 181 mbar pressure, the specimens could be improved (**Supplementary Fig. 6b** and **c**). When the crystal suspension was very viscous, such as containing 35% PEG 6000, high pressure is necessary in order to obtain usable grids (**Supplementary Fig. 6d** and **h**). In reality, the pressure could be further increased to reduce the vitrified ice thickness for the cases as shown in **Supplementary Fig. 6c** and **d**. In conclusion, grids with large hole sizes and high pressure are recommended for viscous crystal suspensions. In cases of crystals grown in extremely viscous buffers such as LCPs, it is difficult to obtain usable EM grids by Preassis alone, a combination with other methods may be necessary, e.g. adding detergents, oils, or lipase(Zhu *et al*., 2020) to decrease the viscosity of the LCP.

Even if a good or usable specimen is obtained, finding ideal grid squares and good crystals for data collection is also important. This is because the thickness distribution of the amorphous ice layer is often not homogeneous across the grid, sometimes is not uniform within the same grid square. For crystals grown in non-viscous mother liquors, ideal grid squares and crystals have the following features: 1) grid squares with clear and sharp edges when viewed in low magnification (**Fig. 3a**), indicating the vitrified ice is very thin, 2) crystals found in these squares with blurred edges on the carbon film, indicating the crystals are still hydrated, and 3) crystals hanging over the empty holes with relatively clear edges, indicating the ice is very thin. It is desirable to collect MicroED data from these types of crystals, and at the regions where they are hanging over the holes. In these cases, carbon background is avoided, the ice thickness is minimized, and the crystals are usually still hydrated. A representative example including images and diffraction patterns is shown in **Fig. 3a**. When the buffer is viscous (e.g. 44% PEG 400 and 30% PEG 4000), the grid squares no longer show sharp edges and transparent areas are reduced compared to those prepared under non-viscous conditions, as shown in the low magnification images in **Fig. 3b** and **c**. Under such circumstances, the ideal crystals for MicroED data collection are still those hanging over the holes (crystal images and ED patterns in **Fig. 3b** and **c**) despite the surrounding vitrified ice. Furthermore, for both viscous and nonviscous cases, most holes in the optimized grid squares are empty without vitrified ice layer except for the holes with crystals nearby or on top of the holes.

**Fig 3.**
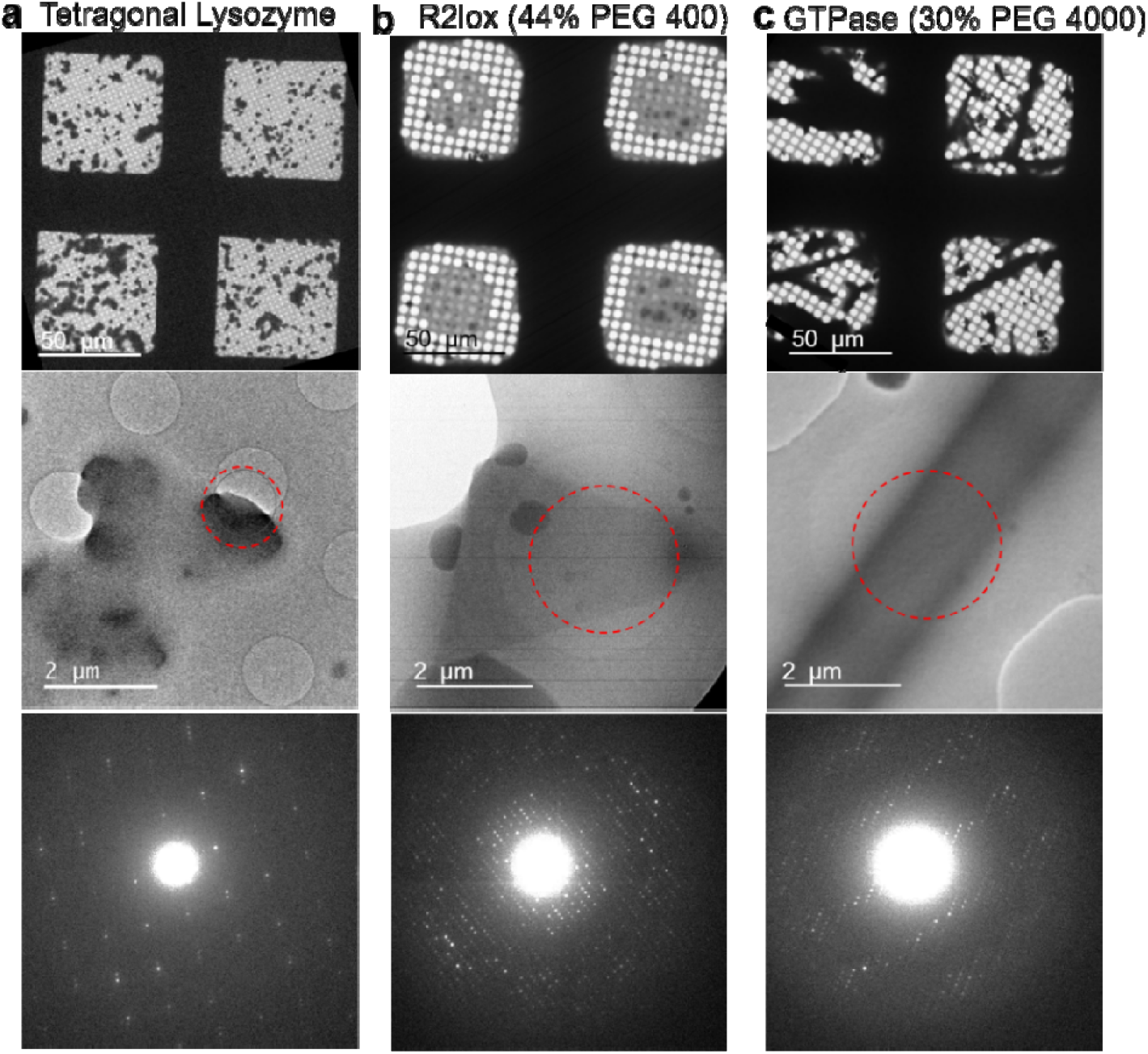
Examples of specimens prepared successfully by Preassis from non-viscous to viscous buffer conditions. **a**, MicroED specimens prepared from tetragonal lysozyme crystal sample under non-viscous condition (without high molecular weight polymer in the buffer) using grid R 1.2/1.3 and pressure of 37.2 mbar. **c**-**d**, MicroED specimens prepared from R2lox and GTPase crystal samples under viscous condition with 44% PEG 400 and 35% PEG 4000, respectively. The pressure used for R2lox specimen is 30.7 mbar, and GTPase is 181 mbar. Grids R 3.5/1 were used for both experiments.

In conclusion, Preassis is a simple and promising method for preparing MicroED specimens. Our results demonstrate that the ice thickness can be adjusted by tuning pressure and selecting grids with different hole sizes. More importantly, Preassis allows the preparation of MicroED specimens for crystals grown in high viscous media and keeps a relatively high density of crystals on the EM grid. Here we provide a guideline on how to select parameters for specimen preparation of a new protein crystal sample using Preassis. Common features of good grid squares and crystals for high-quality MicroED data collection are discussed, which will be important for future automation of data collection and high-throughput structure determination by MicroED. Preassis is simple and can be easily implemented in all cryo-EM labs. New implementations of Preassis, such as developing automation and adding an environmental chamber, may improve the throughput and reproducibility of MicroED specimen preparation. We believe that this method will significantly widen the application of MicroED.

## Methods

### Protein crystallization

#### Needle-shaped orthorhombic lysozyme crystal

Hen egg-white orthorhombic lysozyme microcrystal sample was produced as described by Xu et al.^23^. In brief, the protein was crystallized using the hanging-drop vapour diffusion method where 2 µl of an 8 mg/ml protein solution in 10 mM Tris-HCl pH 8.0 was mixed with 2 µl of reservoir solution consisting of 1 M potassium nitrate, 0.1 M sodium acetate trihydrate pH 3.4, to produce thin fibrous needle-shaped crystal clusters within 48 h at 21 °C.

#### Tetragonal lysozyme crystal

Hen egg-white tetragonal lysozyme microcrystals were grown as described previously(Barends *et al*., 2014; Falkner *et al*., 2005) omitting the cross-linking step and stored in 8% (w/v) NaCl, 0.1 M sodium acetate buffer, pH 4.0. The original crystal suspension was further diluted by 500 times using the same buffer for MicroED specimen preparation.

#### Sulfolobus acidocaldarius R2-like ligand-binding oxidase (SaR2lox)

Crystal sample of the *Sa*R2lox protein was produced as described by Xu et al.(Xu *et al*., 2019). In brief, the protein was crystallized using the hanging-drop vapour diffusion method where a volume of 2 μl of an 8 mg/ml protein solution was mixed with 2 μl of reservoir solution consisting of 44% (v/v) PEG 400, 0.2 M lithium sulphate and 0.1 M sodium acetate pH 3.4. Plate-like crystals grew within 48 h at 21 °C.

#### GTPase crystal

Crystallization of human dynamin I BSE-GTPase construct was performed using sitting-drop vapour-diffusion at 20 °C. Screening of crystallization conditions was carried out with JBScreen Classic 1 and 2 (Jena Bioscience, Jena, Germany). 2 µl of 9 mg/ml protein solution in 20 mM HEPES-NaOH pH 7.5 and 150 mM NaCl buffer were mixed with an equal volume of reservoir solution and equilibrated against 0.5 ml of reservoir solution. The reservoir solution contained 30% (w/v) PEG 4000, making the solution highly viscous. The initial tiny needle-like crystals grew to micrometer size within a day and are suitable for MicroED experiments.

### Specimen preparation by Preassis

#### Specimens of orthorhombic and tetragonal lysozyme crystals

Quantifoil grids R 1.2/1.3 and R 3.5/1 were used in the specimen preparation. The grids were glow discharged with 20 mA current for 60 s by PELCO easiGlow^™^ 9100. A droplet of 3 μl (the same for all other specimen preparations in this work) of crystal suspensions was applied to a glow-discharged grid. The pressures used were 17.2 mbar and 27.7 mbar for the orthorhombic lysozyme sample, 0 mbar and 37.2 mbar for the tetragonal lysozyme sample. The time from applying the sample on the grid to manually plunge-freezing was approximately 5 s for both specimen preparations. Room temperature and humidity were used for all specimen preparations by Preassis.

#### Specimens of SaR2lox and GTPase crystals

Quantifoil grids R 3.5/1 were used for both specimen preparations. The glow-discharge conditions were the same as above. The pressure and time used for *Sa*R2lox and GTPase specimens were 30.7 mbar, 5 s, and 181 mbar, 10 s, respectively.

#### Specimens of samples with different viscosities (concentration of PEG (w/v))

Since it is difficult to get the same type of protein crystals grown in mother liquors of different viscosities, micro-sized inorganic crystals (ZSM-5(Liu *et al*., 2012)) mixed with different concentrations (15%, 25% and 35%) of PEG 6000 and 40% PEG 400 were used. Quantifoil grids R1.2/1.3, R2/1, and R3.5/1 were used in these experiments and were glow discharged as above. For the specimens with PEG 6000, the pressure was either 0 mbar or 181 mbar, and the time from applying the sample on the grid to manually plunge-freezing was ca 10 s. For the specimens with 40% PEG 400, the pressure was 78 mbar and the time was ca 5 s.

### Specimen preparation by Vitrobot Mark IV (Thermo Fisher Scientific)

#### Specimens of R2lox crystal

A droplet of 3 μl of R2lox crystal suspension was applied to a glow-discharged (60 s, 20 mA, PELCO easiGlow) Quantifoil grid R3.5/1. The operation parameters of the Vitrobot Mark IV were 4 °C, 100% humidity, 10 s blotting time, two layers of blotting papers on each pad, and 16 blotting force.

#### Specimens of orthorhombic lysozyme crystal

A droplet of 3 μl of lysozyme crystal suspension was applied to a glow-discharged (60 s, 20 mA, PELCO easiGlow) Quantifoil grid (R 3.5/1). The operation parameters of the Vitrobot Mark IV (Thermo Fisher Scientific) were 4 °C, 100% humidity, single blot, 5 s blotting time, 1 blotting paper on each pad, and −6 blotting force.

#### Specimens of crystal suspension with 40% PEG 400

The sample was prepared by mixing ZSM-5 microcrystals with 40% (w/v) PEG 400. Quantifoil grids R2/1 were used and glow-discharged as above. A 3 μl droplet of the crystal suspension was applied to a glow-discharged grid. The operation parameters of the Vitrobot were room temperature, single blot, 5 s blotting time, 1 blotting paper on each pad, and 0 blotting force. The humidity used for these specimens was either room humidity (35-45%) or 100% humidity.

### TEM image collection

TEM images were collected on a JEOL JEM-2100LaB_6_ TEM equipped with an Orius detector. All the images were collected at 200 kV under cryogenic condition using a Gatan 914 cryo-transfer holder. Because of the lens distortion at ultra-low magnification, the images of the grid maps are distorted, especially at the edges.

### Electron diffraction data collection

Selected area electron diffraction patterns were collected under cryogenic conditions using a Gatan 914 cryo-transfer holder on a JEOL JEM-2100LaB_6_ TEM operated at 200 kV, and recorded by a fast Timepix hybrid pixel detector (Amsterdam Scientific Instruments). The conditions used to collect data were: spot size 3, cameral length 80 cm / 100 cm, and exposure time 1 s / 2 s per frame.

## Acknowledgments

The project is supported by the Knut and Alice Wallenberg Foundation (2012.0112 and 2018.0237, X.Z.; 2017-04018, M.H.), the Swedish Research Council (2017-04018, M.H.; 2017-05333, H.X.; 2019-00815, X.Z.) and the Science for Life Laboratory through the pilot project grant Electron Nanocrystallography, and MicroED@SciLifeLab. The authors acknowledge the Cryo-EM Swedish National Facility jointly funded by the Knut and Alice Wallenberg, Family Erling Persson and Kempe Foundations, SciLifeLab, Stockholm University, and Umeå University. The authors also thank I. Schlichting at Max-Plank Institute for Medical Research for providing the tetragonal lysozyme sample.

## Author contributions

J.Z. contributed to method development, design and tests of the Preassis setup, specimen preparation, TEM and MicroED data collection, data processing and analysis, initial manuscript preparation and figure preparation. H.X. contributed to project design, method development, specimen preparation, MicroED data collection, and manuscript preparation. M.C. contributed to the project discussion. H.L., H.T. and M.H. prepared protein microcrystals and contributed to the project discussion. X.Z. contributed to project design, the conception of Preassis, and manuscript preparation. All authors contributed to the method discussions and manuscript revisions. H.X. and X.Z. led the project and finalized the manuscript.

## Supplementary protocol

Preassis is widely applicable for MicroED specimen preparation, including crystals grown in low-viscous buffer conditions to highly-viscous PEG-rich crystallization conditions. It must be noted that different crystal samples grown in different conditions and with different sizes or shapes behave differently. The parameters for the method described here may need to be fine tuned accordingly. Furthermore, the pressure range also depends on the size of the Büchner flask and the type of pump machine. Therefore, the parameters applied here for different types of protein crystal samples may need to be adjusted for a new setup. The specimen preparation comprises the following steps:

a. providing an EM grid;
b. applying a sample suspension onto the EM grid;
c. providing a pressure gradient through the EM grid in order to pull a portion of the liquid through the grid while removing the excess liquid with a filter paper on the other side of the grid;
d. plunging the specimen into a cryogenic bath.

Details of the setup and the specimen preparation procedure are described below.

### Equipment

- Micropipette and tips, 0.5-10 μL (Eppendorf Research plus variable micropipette, #3116000015 or similar; Eppendorf tips 0.1-10 μL, # Z741098-960EA or similar)
- Tweezers (Dumont Tweezer, style5 #72705-01or similar)
- Filter paper (Munktell, #110067 or similar)
- Quantifoil holey carbon grids (R 1.2/1.3, R 2.0/1, R 3.5/1, R2/2, or similar)
- Glow-discharge (PELCO easiGlow^™^ 9100 or similar)
- Cryo grid box (CGB4-1 SWISSCI Cryo grid box or similar)
- FEI coolant container
- Ethane and liquid nitrogen
- Vacuum aspirator or pump machine (PC 3001 VARIO)
- Polymer hose (hose, PVC-/latex-, 6 × 9 mm)
- Pressure meter (PELCO^R^ 2245 Minl Hot Vac. if a vacuum aspirator is used)
- Büchner flask (GLASSCO 500 mL, or similar)

### Procedure

1. Prepare liquid ethane using the FEI coolant container.
2. Glow discharge the grids (current 20 mA, glow 60 s, hold 10 s; can be changed if necessary). The hole size of the grid should be slightly smaller than the crystal size if possible. For very viscous samples, grids with large hole sizes are recommended, such as Quantifoil R 3.5/1 grids.
3. Put a filter paper on top of the flask.
4. Use a pair of tweezers to pick up a glow discharged grid and put it onto the center of the filter paper with the carbon side facing up.
5. Turn on the tap (if a vacuum aspirator is used) or start the vacuum pump. If a vacuum aspirator is used, a pressure meter needs to be installed to measure the pressure. For non-viscous samples, typical pressures used for MicroED specimen preparation are around 20 mbar if a Quantifoil R1.2/1.3 is used (the pressure is not always necessary when a grid with larger hole size is used, e.g Quantifoil R2/1 and R3.5/1). For viscous samples, a pressure of 70 mbar or higher may be needed even for grids with large hole sizes, e.g. Quantifoil R3.5/1.
6. Use a micropipette to take 3 μl of the sample, apply it onto the grid. The droplet will be dispersed through the grid immediately. Pick the grid up and plunge freeze it within 5 to 10 s. For very viscous samples, one can decrease the sample volume to 1 or 2 μl, use a grid with large hole size (e.g Quantifoil R 3.5/1), increase the pressure strength, or wait for a slightly longer time before picking up the grid.
7. Cryotransfer the grid into a cryo-grid box.

In this setup, a vacuum aspirator with water-flow or a vacuum pump machine (recommended) can be used to produce the pressure/suction. In principle, the suction time can also influence the thickness of the vitrified ice. However, the time is less controllable compare with the other two parameters at the current stage. We noticed that the hole size and pressure play more important roles on the ice thickness. We admit that there is a small deviation between specimens prepared using the same parameters, which may result from the manual picking, time controlling, and also the contact between the grid and filter paper. If we handle the process properly, the deviation is small and will not affect the overall results. In the future, new implementations, such as automation development and humidity chamber, may improve the throughput and reproducibility of the specimen preparation by Preassis.

## Supplementary Figures

**Supplementary Fig. 1.**
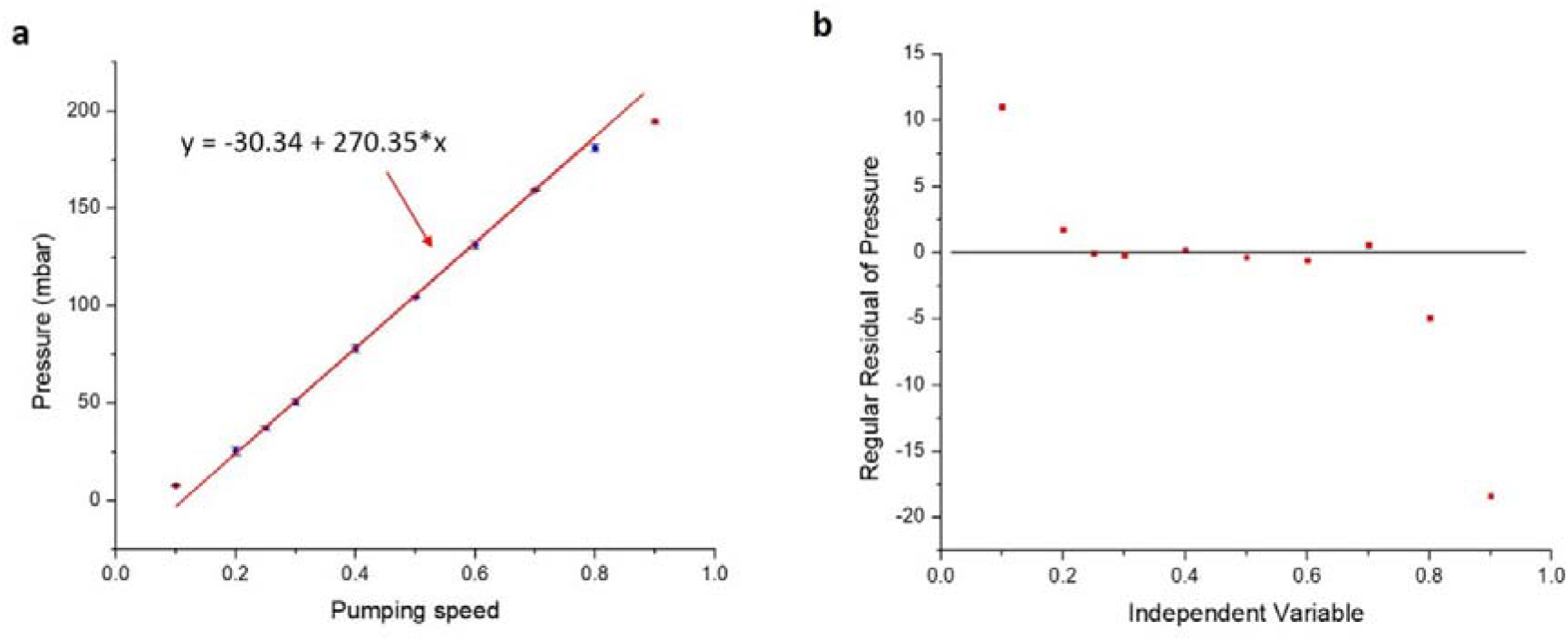
Relationship between the pumping speed and the resulting pressure of the pumping machine (PC 3001 VARIO). a-b, The pressure is almost linearly related to the pumping speed in the region where the pumping speed is between 20% and 80%. The regular residual of the pressure in **b** is defined by the difference between the predicted value by *y* = −30.34+ 270.35\**x* in **a** and the experimental value.

**Supplementary Fig. 2.**
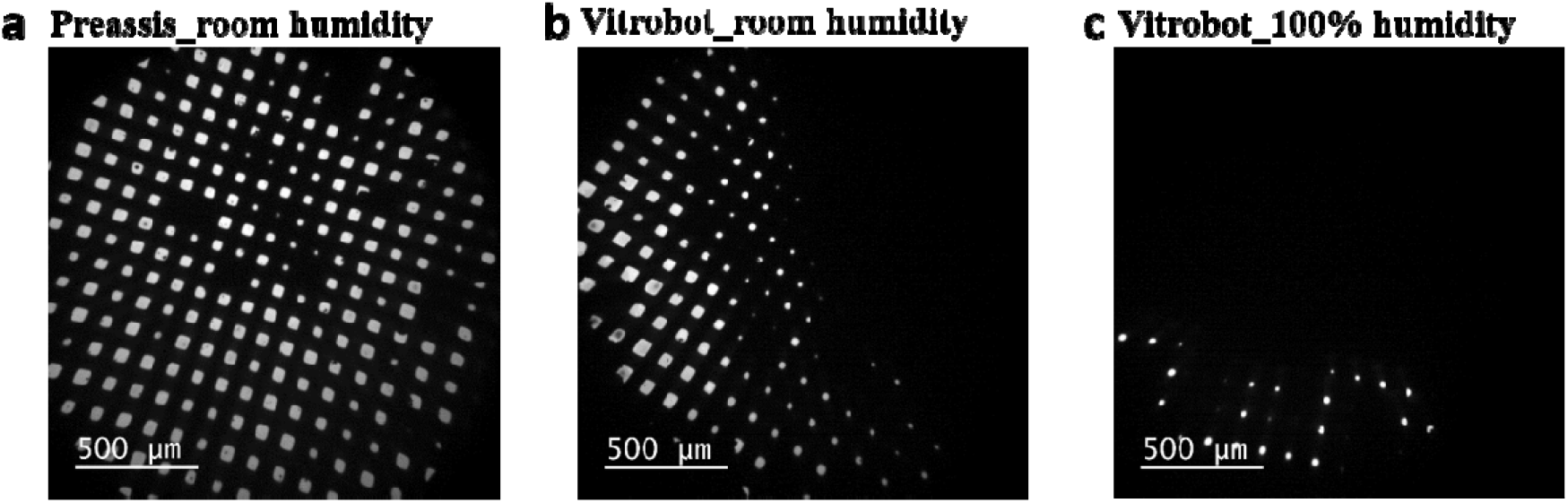
Comparison of EM grids prepared by Preassis and Vitrobot under different humidity using a mixture of ZSM-5 microcrystals with 40% PEG 400. **a**, MicroED specimens prepared by Preassis with pressure of 78 mbar (5 s) and **b**, Vitrobot with 0 blotting force (5 s). These two specimens were prepared under room humidity (35% −45%) and temperature (ca 20 °C). **c**, Specimen prepared by Vitrobot under 100% humidity and room temperature. This comparison illustrates that Preassis is more efficient in removing liquid comparing with Vitrobot. High humidity reduces the liquid removal efficiency as caused by the decrease of the liquid adsorption ability of filter paper. R 2/1 grids were used for these three experiments. Because of the lens distortion at such low magnification, the images are distorted especially at the edges.

**Supplementary Fig. 3.**
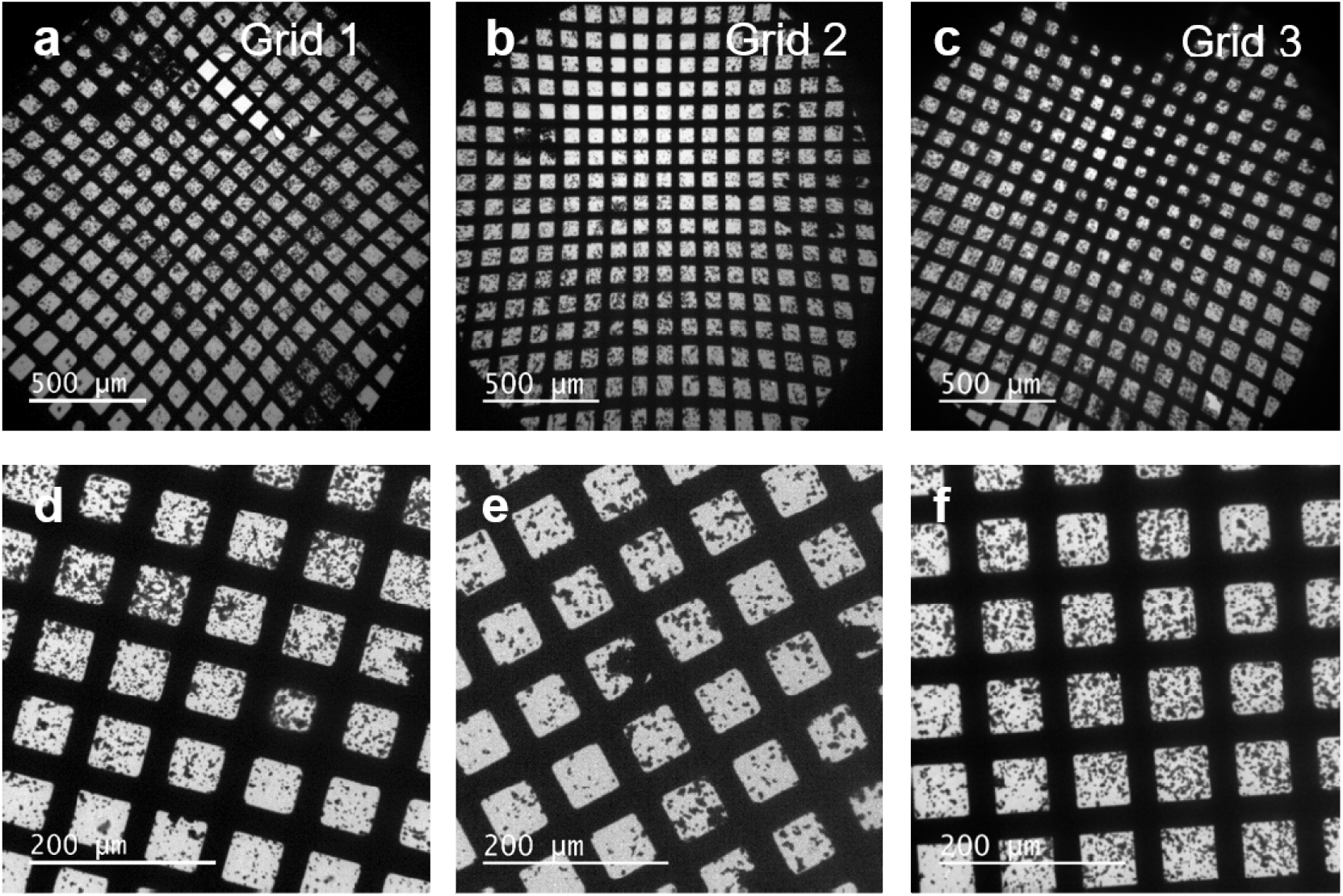
Reproducibility of specimen preparation by Preassis using tetragonal lysozyme crystal sample. **a-c**, Low magnification TEM images of MicroED specimens prepared by Preassis using the same experimental parameters: 3 μl droplet, Quantifoil grid R 1.2/1.3, 37.2 mbar pressure, suction time ca 5 s, and room temperature (ca 20 °C) and humidity (35% −45%). Because of the lens distortion at such low magnification, the images are distorted especially at the edges. **d-f**, TEM images taken from those three repeated grids showing the local ice conditions.

**Supplementary Fig. 4.**
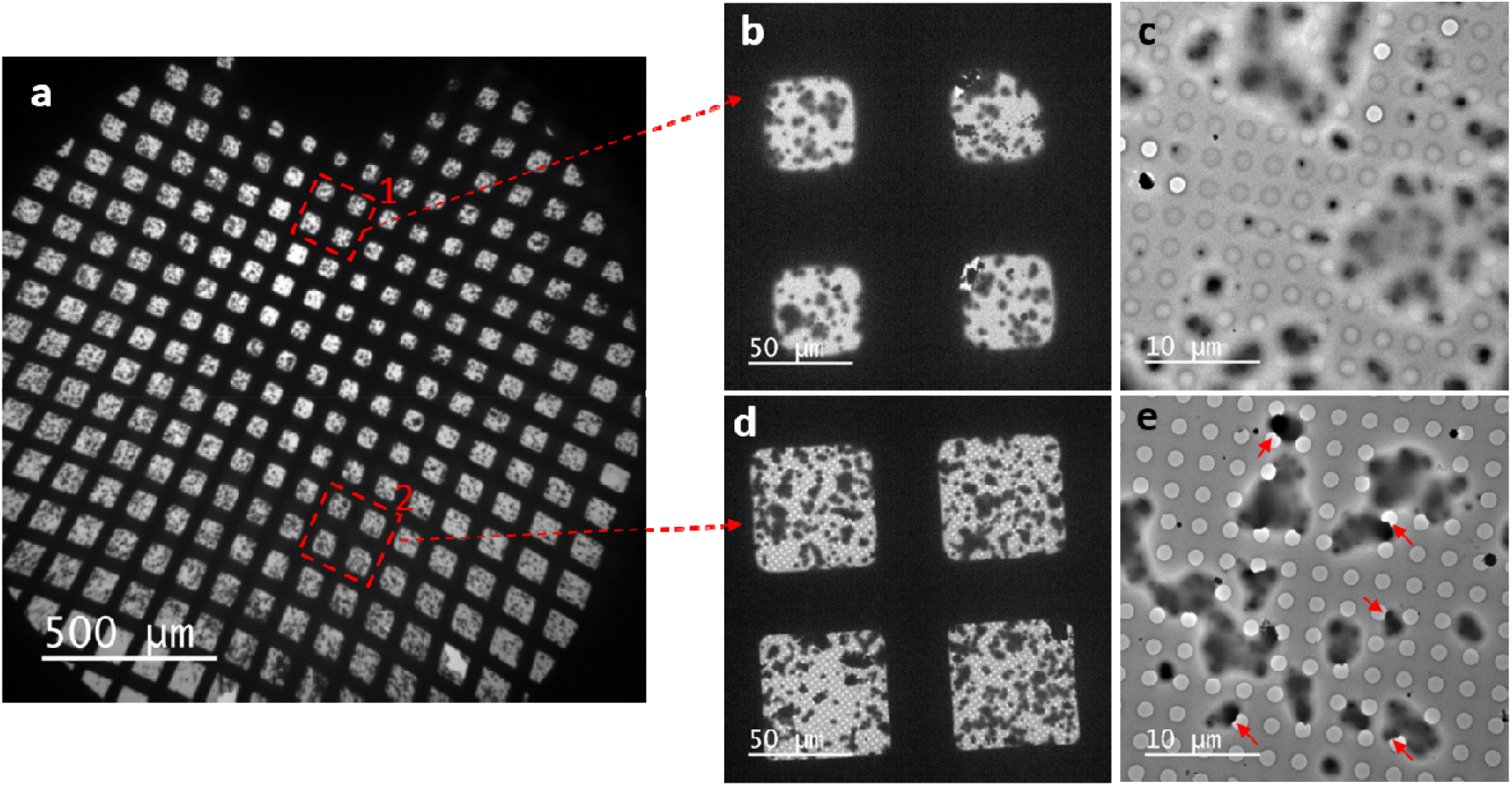
Ice thickness distribution across the grid prepared by Preassis from the tetragonal lysozyme crystal sample. **a**, Low magnification image showing the inhomogeneous distribution of ice thickness on the entire TEM grid. Because of the lens distortion at such low magnification, the image is distorted especially at the edges. **b-c** and **d-e**, Two representative examples of areas with relatively thick (**b-c**) and thin (**d-e**) ice layers. The crystals marked by red arrows in image **e** are suitable for MicroED data collection.

**Supplementary Figure 5.**
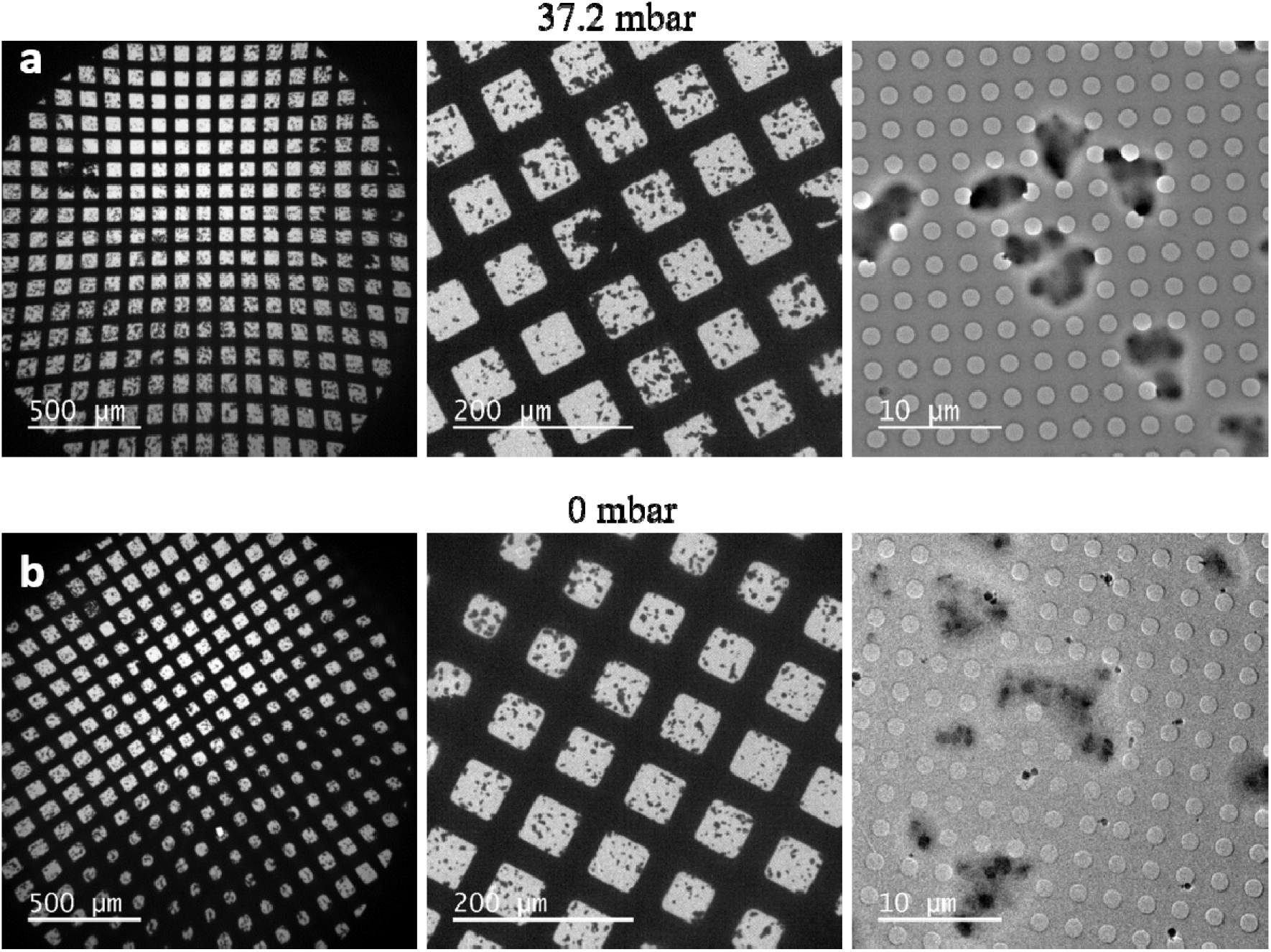
Influence of pressure on MicroED specimen prepared from a non-viscous buffer by Preassis using a tetragonal lysozyme crystal sample. **a** and **b**, Low magnification images and high magnification images taken from the specimens prepared with pressure of 37.2 mbar and 0 mbar, respectively. For these specimens, tetragonal lysozyme crystal sample and Quantifoil grid R 1.2/1.3 were used. Because of the lens distortion at such low magnification, the image is distorted especially at the edges.

**Supplementary Figure 6.**
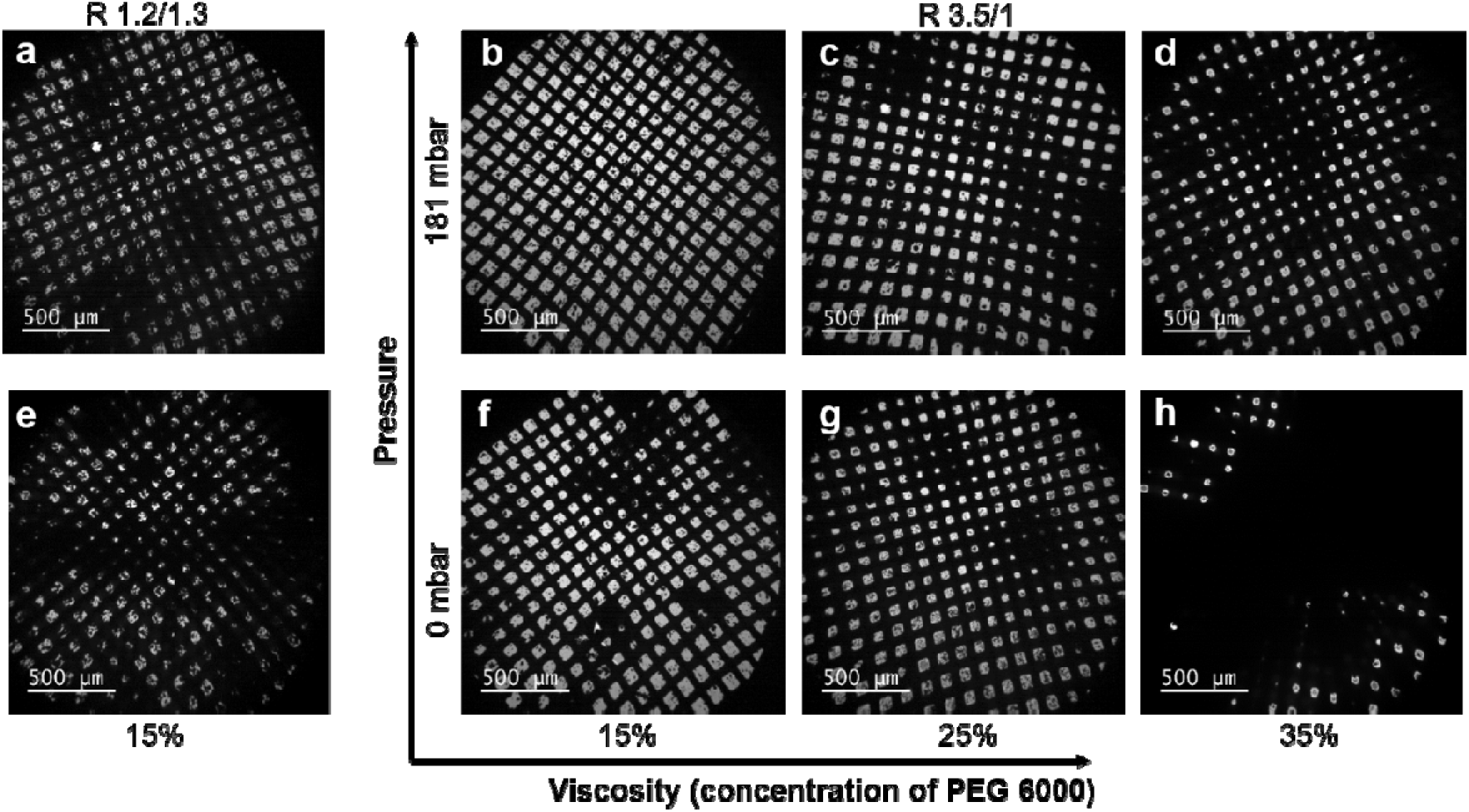
Influence of pressure and hole size on MicroED specimens prepared by Preassis using crystal suspensions with different viscosity. **a** and **e**, Low magnification TEM images taken from the specimens prepared from a crystal suspension with 15% PEG 6000 using Quantifoil grids R 1.2/1.3 with 181 and 0 mbar pressure, respectively. **b**-**d** and **f**-**h**, Low magnification TEM images taken form the specimens prepared by Preassis from crystal suspensions with different viscosity (15%, 25%, and 35% PEG 6000) using Quantifoil grids R 3.5/1 at 181 and 0 mbar pressure, respectively. The crystal suspensions with different viscosities were made by mixing ZSM-5 microcrystals with different concentration of PEG 6000.

## Reference

Abdallah, B. G., Roy-Chowdhury, S., Fromme, R., Fromme, P. & Ros, A. (2016). Cryst. Growth Des. 16, 2074–2082.

Barends, T. R. M., Foucar, L., Botha, S., Doak, R. B., Shoeman, R. L., Nass, K., Koglin, J. E., Williams, G. J., Boutet, S., Messerschmidt, M. & Schlichting, I. (2014). Nature. 505, 244–247.

Beale, E. V., Waterman, D. G., Hecksel, C., van Rooyen, J., Gilchrist, J. B., Parkhurst, J. M., de Haas, F., Buijsse, B., Evans, G. & Zhang, P. (2020). Front. Mol. Biosci. 7,.

Caffrey, M. (2015). Acta Crystallogr. Sect. F Struct. Biol. Commun. 71, 3–18.

Clabbers, M. T. B. & Xu, H. (2020). Drug Discov. Today Technol.

de la Cruz, M. J., Hattne, J., Shi, D., Seidler, P., Rodriguez, J., Reyes, F. E., Sawaya, M. R., Cascio, D., Weiss, S. C., Kim, S. K., Hinck, C. S., Hinck, A. P., Calero, G., Eisenberg, D. & Gonen, T. (2017). Nat. Methods. 14, 399–402.

Dubochet, J. & McDowall, A. W. (1981). J. Microsc. 124, 3–4.

Duyvesteyn, H. M. E., Kotecha, A., Ginn, H. M., Hecksel, C. W., Beale, E. V., de Haas, F., Evans, G., Zhang, P., Chiu, W. & Stuart, D. I. (2018). Proc. Natl. Acad. Sci. 115, 9569–9573.

Falkner, J. C., Al-Somali, A. M., Jamison, J. A., Zhang, J., Adrianse, S. L., Simpson, R. L., Calabretta, M. K., Radding, W., Phillips, G. N. & Colvin, V. L. (2005). Chem. Mater. 17, 2679–2686.

Gemmi, M., La Placa, M. G. I., Galanis, A. S., Rauch, E. F. & Nicolopoulos, S. (2015). J. Appl. Crystallogr. 48, 718–727.

Li, X., Zhang, S., Zhang, J. & Sun, F. (2018). Biophys. Rep. 4, 339–347.

Liu, F., Willhammar, T., Wang, L., Zhu, L., Sun, Q., Meng, X., Carrillo-Cabrera, W., Zou, X. & Xiao, F.-S. (2012). J. Am. Chem. Soc. 134, 4557–4560.

Martynowycz, M. W., Shiriaeva, A., Ge, X., Hattne, J., Nannenga, B. L., Cherezov, V. & Gonen, T. (2020). BioRxiv. 2020.09.27.316109.

Martynowycz, M. W., Zhao, W., Hattne, J., Jensen, G. J. & Gonen, T. (2019). Structure. 27, 1594–1600.e2.

McPherson, A. & Gavira, J. A. (2014). Acta Crystallogr. Sect. F Struct. Biol. Commun. 70, 2–20.

Nannenga, B. L. & Gonen, T. (2019). Nat. Methods. 16, 369–379.

Nannenga, B. L., Shi, D., Leslie, A. G. W. & Gonen, T. (2014). Nat. Methods. 11, 927.

Nederlof, I., van Genderen, E., Li, Y.-W. & Abrahams, J. P. (2013). Acta Crystallogr. D Biol. Crystallogr. 69, 1223–1230.

Polovinkin, V., Khakurel, K., Babiak, M., Angelov, B., Schneider, B., Dohnalek, J., Andreasson, J. & Hajdu, J. (2020). J. Appl. Crystallogr. 53, 1416–1424.

Russo Krauss, I., Merlino, A., Vergara, A. & Sica, F. (2013). Int. J. Mol. Sci. 14, 11643–11691.

Shi, D., Nannenga, B. L., de la Cruz, M. J., Liu, J., Sawtelle, S., Calero, G., Reyes, F. E., Hattne, J. & Gonen, T. (2016a). Nat. Protoc. 11, 895.

Shi, D., Nannenga, B. L., Iadanza, M. G. & Gonen, T. (2013). ELife. 2,.

Xu, H., Lebrette, H., Clabbers, M. T. B., Zhao, J., Griese, J. J., Zou, X. & Högbom, M. (2019). Sci. Adv. 5, eaax4621.

Xu, H., Lebrette, H., Yang, T., Srinivas, V., Hovmöller, S., Högbom, M. & Zou, X. (2018a). Structure. 26, 667–675.e3.

Zhou, H., Luo, Z. & Li, X. (2019). J. Struct. Biol. 205, 59–64.

Zhu, L., Bu, G., Jing, L., Shi, D., Lee, M.-Y., Gonen, T., Liu, W. & Nannenga, B. L. (2020). Structure.

